# Immunoinformatic approach to design CTL epitope based chimeric vaccine targeting multiple serotypes of dengue virus

**DOI:** 10.1101/2024.01.15.575641

**Authors:** Nilanshu Manocha, Prashant Kumar, Madhu Khanna

**Author notes:** These authors contributed equally (Nilanshu Manocha, Prashant Kumar).

## Abstract

The dengue outbreak is one of the serious global public health concerns. The World Health Organization reported 3,80,171 cases and 113 deaths this year till March 2023, and the rate of infection is expected to increase in vulnerable parts of the world. The development of vaccines is the best approach to managing infectious diseases. All the approved vaccines against dengue are based on live-attenuated virus but they have been questioned for their effectiveness in some population categories. Additionally, random occurrence of four closely related serotypes of dengue virus in humans leading to antibody-dependent enhancement of the disease is yet another cause of vaccine ineffectiveness. Therefore, development of a therapeutic subunit-vaccine based on epitopes from all four serotypes may be expected to provide effective cross-protective cellular immunity. Towards this end, we designed a multi-epitope chimeric immunogen using envelop protein of dengue virus. The MHC-I binding T-cell epitopes were predicted based on their immunogenicity, allergenicity and antigenicity. NetMHCpan-EL4.1 prediction method was used to determine the binding ability of the epitopes with HLA alleles with population coverage of over 99%. The five most potent epitopes based on their immunogenicity, population coverage and prediction scores were selected for each serotype and a multi-epitope polypeptide was generated by merging peptides with AAY linker. The polypeptide was predicted to be an antigen and a non-allergen with a stable tertiary structure retaining a half-life of 4.4 hours in mammalian system. The polypeptide has the potential to elicit effective cellular immune response against all the dengue virus serotypes.

## Introduction

Dengue virus (DENV) is by far one of the most prevalent mosquito-borne arboviruses affecting humans worldwide. Dengue infection is commonly transmitted by the bite of an infected female *Aedes* spp. mosquito, which breeds in peridomestic and tropical/ subtropical environments (*Aedes aegypti*). It is predicted that future climate trends will increase the risk of establishing another less prevalent *Aedes albopictus* mosquito in northern Europe due to wetter and warmer conditions (Mercier et al., 2022). Four closely related serotypes of the virus (DENV-1, DENV-2, DENV-3, and DENV-4) cause dengue disease. The immunity against cross-serotype infections in an average young adult is only partial and temporary. Partial immunity acts as a catalyst for disease severity (Sarker et al., 2023). An estimated 3.9 billion people are at risk of infection, of which 96 million people manifest clinically with severe complications such as dengue haemorrhagic fever (DHF) or dengue shock syndrome (DSS) (Bhatt et al., 2013). Targeting our immune system has the potential to overcome the disease burden. Although there are a few dengue vaccines in the pipeline, only one tetravalent live attenuated vaccine (Dengvaxia; Sanofi Pasteur) is approved by the regulatory authorities in nearly 20 countries and has been shown efficacious only in seropositive individuals (Amorim et al., 2022).

Several studies show the promise of either natural or novel synthetic peptides in activating T cells to combat viral infections (Tian et al., 2019; Adikari et al., 2020). In this study, we have initiated the development of peptide-based candidates as potential immune activators against DENV. Multi-disciplinary strategies are required in conjunction to design and develop such candidates. The foremost approach is to optimise immunodominant peptide selection using advanced immunoinformatics tools. Such tools run sophisticated and well-tuned machine learning algorithms, enabling analysts to swiftly predict potential immunogenic epitopes (Ruth et al., 2015). The Immune Epitope Database (IEDB) is one such freely available resource (www.iedb.org), which index an extensive collection of experimental data on T cell epitopes studied in humans and other clinically relevant animal species (Vita et al. 2019). Currently, the database has collected over 1.5 million epitopes and more than 6.5 million T cell, B cell, MHC binding, and MHC ligand elution assays. The IEDB repository has refined the experimental data curated from over 23,500 referenced scientific studies in infectious diseases, allergy, autoimmunity, and transplantation (Fleri et al., 2017). Using IEDB prediction and analysis tools, we can query the protein sequences for known epitopes and their immunogenicity. The algorithms of these tools analyse peptide sequence patterns to determine their binding affinity to MHC molecules based on the structural similarity and molecular docking data.

In the last decade, evolution in data sciences and machine-learning models has significantly improved the prediction accuracy of such tools and meta-analyses. Here we have performed an intensive and comprehensive *in silico* study to select major histocompatibility complex (MHC) class I binding epitopes and linked them to design a vaccine construct targeting all the serotypes of DENV. Since the envelope (E) protein of DENV is a critical determinant of infectivity, we predicted the best MHC class I-binding epitopes from the E protein. The vaccine construct was predicted to have high immunogenicity with high population coverage and with a potential to generate effective immune response. Although the *in-silico* analysis shows the potential of our vaccine construct, its potentiality needs *in vitro* and *in vivo* experimental validation.

## Methods

### DENV Protein Sequence Retrieval, Conserved Domains Search and Their Transmembrane Properties

The DENV envelope (E) protein reference sequences were retrieved from Virus Variation Database for each serotype (DENV1: NP_722460.2; DENV2: NP_739583.2; DENV3: YP_001531168.2 and DENV4: NP_740317.1).Further, all known E protein sequences of all DENV serotypes were collected in FASTA format with search criteria of type: standard; disease: any; host: human; region: any; genome: E; isolation: any. Multiple sequence alignment (MSA) of the four serotypes was individually performed through the Clustal Omega algorithm, and conserved domains with a parameter of minimum segment length of 15 amino acids were identified using BioEdit software. Later, the conserved sequences were screened for soluble and membrane parts using the DeepTMHMM webserver (https://dtu.biolib.com/DeepTMHMM), a most widely used alpha-helical transmembrane topology model. DeepTMHMM is based upon a deep learning model, where protein sequence is input and outputs a discriminated intracellular (I), membrane (M) or surface/outside cell/lumen of ER/Golgi/lysosomes (O) regions of the protein (Hallgren et al., 2022).

### MHC-I T Cell Epitopes Prediction

The IEDB-recommended NetMHCPan EL 4.1 tool was used to predict MHC-I binding T cell epitopes. The prediction method assigns binding scores or percentile ranks (PRs) to the predicted peptide based on its affinity to the MHC complex (Reynisson et al., 2020). The PR is calculated by comparing its prediction score to a distribution of prediction scores for a selected MHC, estimated from a set of random natural peptides. The PR value of 1.0 indicates that a predicted sequence corresponds to the top 1% scores obtained from random natural peptides.

In this study, IEDB recommended method was selected, and predictions were performed against 9mer peptides. The reference E protein sequences were used as input, and the frequently occurring HLA (human) and H2 (mouse) class I alleles were selected for the prediction analysis. The threshold for PR < 0.5 was set. Predictions were generated against 75 frequently occurring human MHC-I alleles (including 12 HLA supertypes) and six mouse MHC-I alleles. The peptides were screened for inter- and intra-serotype conservation using the conserved domain search data above for each serotype prediction.

### T Cell Epitopes –MHC I Processing Prediction

The proteasomal cleavage, TAP transport and MHC binding were jointly predicted for all the DENV serotypes using the IEDB MHC-I processing prediction tool(http://tools.iedb.org/processing/) with default settings, which uses the NetMHCpan algorithm. Immuno-proteasome was selected as the proteasome type to predict efficient epitopes in the context of antigen presentation. This tool combines the score of all three parameters and predicts an overall score depicting the inherent potential of input peptides as a T cell epitope (Tenzer et al., 2005). Based on this score, the best candidates were selected for further analysis.

### MHC-I Immunogenicity Prediction

The peptides in the above analysis were screened for their immunogenicity as T-cell epitopes using IEDB online server (http://tools.iedb.org/immunogenicity/). The immunogenicity prediction tool integrates amino acid physiochemical properties and its position within the peptide to determine if T cells can recognise and target a pMHC (Calis et al., 2013). Since the tool is well-validated for 9mer peptides, the prediction was performed with default settings that masked first, second and C-terminus amino acids (Trolle et al., 2016).

### Population Coverage Prediction

The MHC genes show high polymorphism, implying that many different HLA alleles are expressed in different groups or ethnicities inside a population and expressed at different genotypic frequencies (Choo, 2007). Therefore, it is crucial to identify that the predicted MHC-allele-specific T cell epitope(s) should cover the diverse human population as a global therapeutic peptide candidate. Addressing the concern, IEDB provides a population coverage analysis tool (http://tools.iedb.org/population/), where the shortlisted epitope sequences with the corresponding HLA I alleles were submitted. This tool calculates the cumulative percentage of individuals predicted to respond to a given epitope set based on HLA genotypic frequencies and on the basis of MHC binding and/or T cell restriction data (Bui et al., 2006). The allele frequency database (http://www.allelefrequencies.net/) is the most comprehensive database of HLA allele genotypic frequencies for 115 countries and 21 different ethnicities grouped into 16 different geographical regions. For a particular population coverage, the IEDB tool calculates three values: projected population coverage, the average number of epitope hits / HLA combinations recognised by the population, and the minimum number of epitope hits / HLA combinations recognised by 90% of the population (PC90).

### Construction of multi-epitope vaccine construct

Five potent MHC-I T cell epitopes from each dengue serotype were selected to design a chimeric vaccine construct, which was prepared by linking the epitopes with AAY linker. The physicochemical properties of the vaccine construct were determined and validated using Protparam server (Zaib et al., 2022).

### Determination of allergenic and antigenic property of vaccine construct

The vaccine construct is a polypeptide and therefore, the allergenic capacity of the construct was determined using online AllerTOP v2.0 tool (https://www.ddg-pharmfac.net/AllerTOP/). The antigenic nature of the construct was determined using VaxiGen v2.0.

### Prediction and validation of structure of vaccine construct

The secondary structure of the vaccine construct was predicted based on its primary amino acid sequence using Psipred online server (http://bioinf.cs.ucl.ac.uk/psipred/). The tertiary structure was predicted with I-Tasser server (https://zhanglab.ccmb.med.umich.edu/I-TASSER/) which uses sequence-structure-function paradigm. The tertiary structure was further validated using the pictorial database, PDBsum which generated Ramachandran plot to evaluate the structure accuracy.

### Docking of the vaccine construct with Toll-like-receptor 3 (TLR3)

TLR3 is known to restrict the dengue virus infection via induction of type I interferon (Liang et al., 2011). Therefore, we performed a docking analysis to assess if the vaccine construct can be recognized by TLR3 thus leading to secretion of antiviral cytokines. The TLR3 was docked with our vaccine construct using PatchDock server and the docking results so obtained were further refined and rescored using FireDock server (Kumar et al., 2022). The best model was selected, and the molecular structure was visualized and analysed with UCSF chimera.

### Optimisation of the vaccine construct

To enhance the efficiency of cloning of DNA encoding the vaccine construct and accuracy of translated sequence, Java Codon Adaptation Tool (JCat) server (http://www.jcat.de/) was used for codon optimization with respect to human cells. The GC-content and Codon Adaptation Index (CAI) value of the optimized sequence was analysed to predict the expression level of the vaccine construct.

### Prediction of immunogenicity of vaccine construct

The chimeric vaccine construct was assessed using online simulation server, C-ImmSim (http://150.146.2.1/C-IMMSIM/index.php) to predict its immunogenicity. The cellular immune interaction in mammalian system was predicted using Machine Learning which utilized position-specific scoring matrix (PSSM). The prediction was done for three doses of vaccines at four weeks interval with the simulation volume of 50 and 1000 simulation steps. The vaccine construct was injected without any LPS.

## Results

### Retrieval of DENV Protein Sequences and Identification of Conserved Domains

Total of 1,937, 1499, 935 and 370 sequences for the DENV1, DENV2, DENV3 and DENV4 respectively were retrieved from the virus variation database in FASTA format. The Clustal Omega algorithm of MSA generated several conserved sequences for each serotype using SnapGene 6.2.1 (for MSA) and BioEdit 7.7.1 (for conserved domain search) (Table 1). DeepTMHMM data of all DENV E proteins except DENV4 E protein fulfill the criterion of exomembrane characteristics.

**Table 1:**
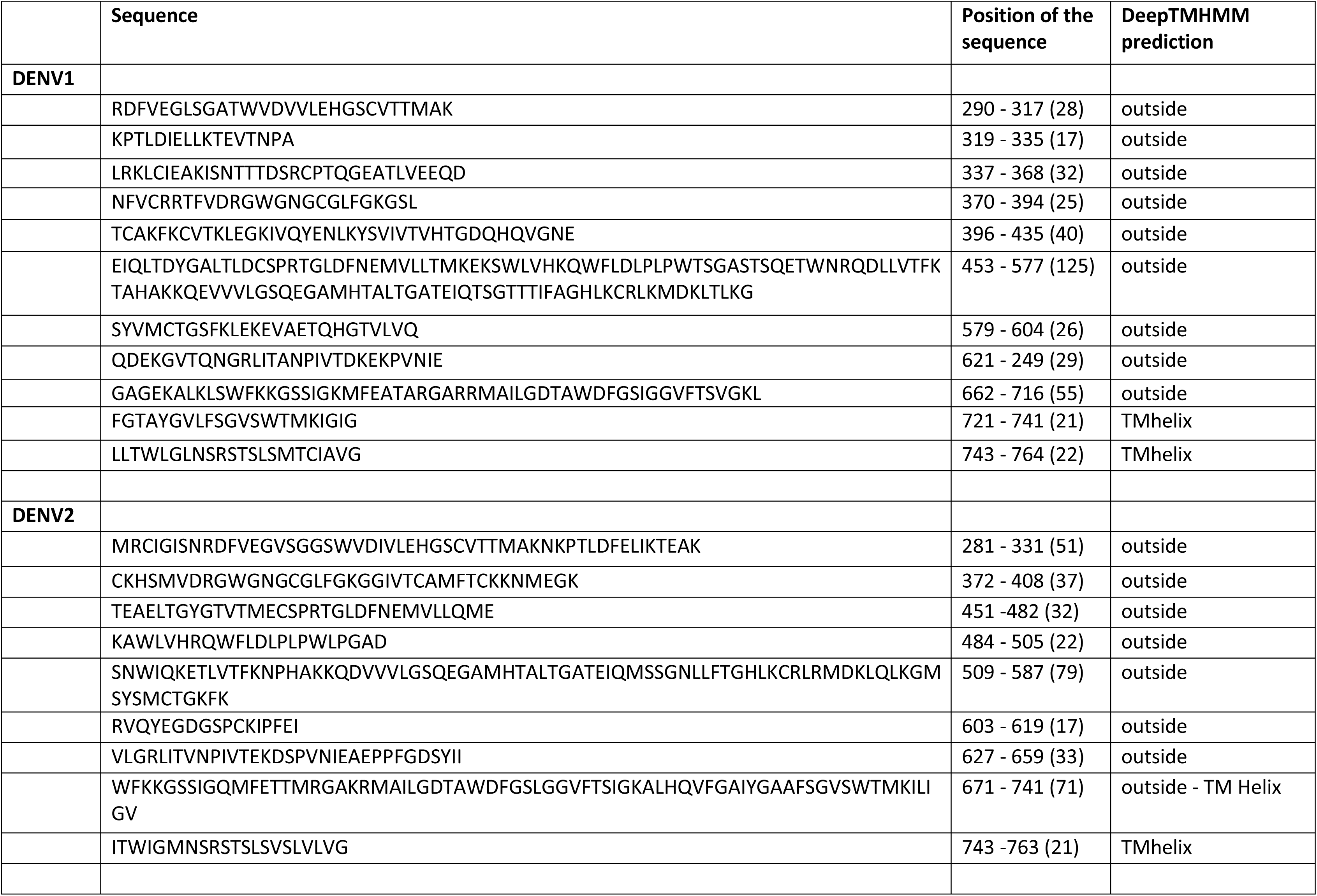
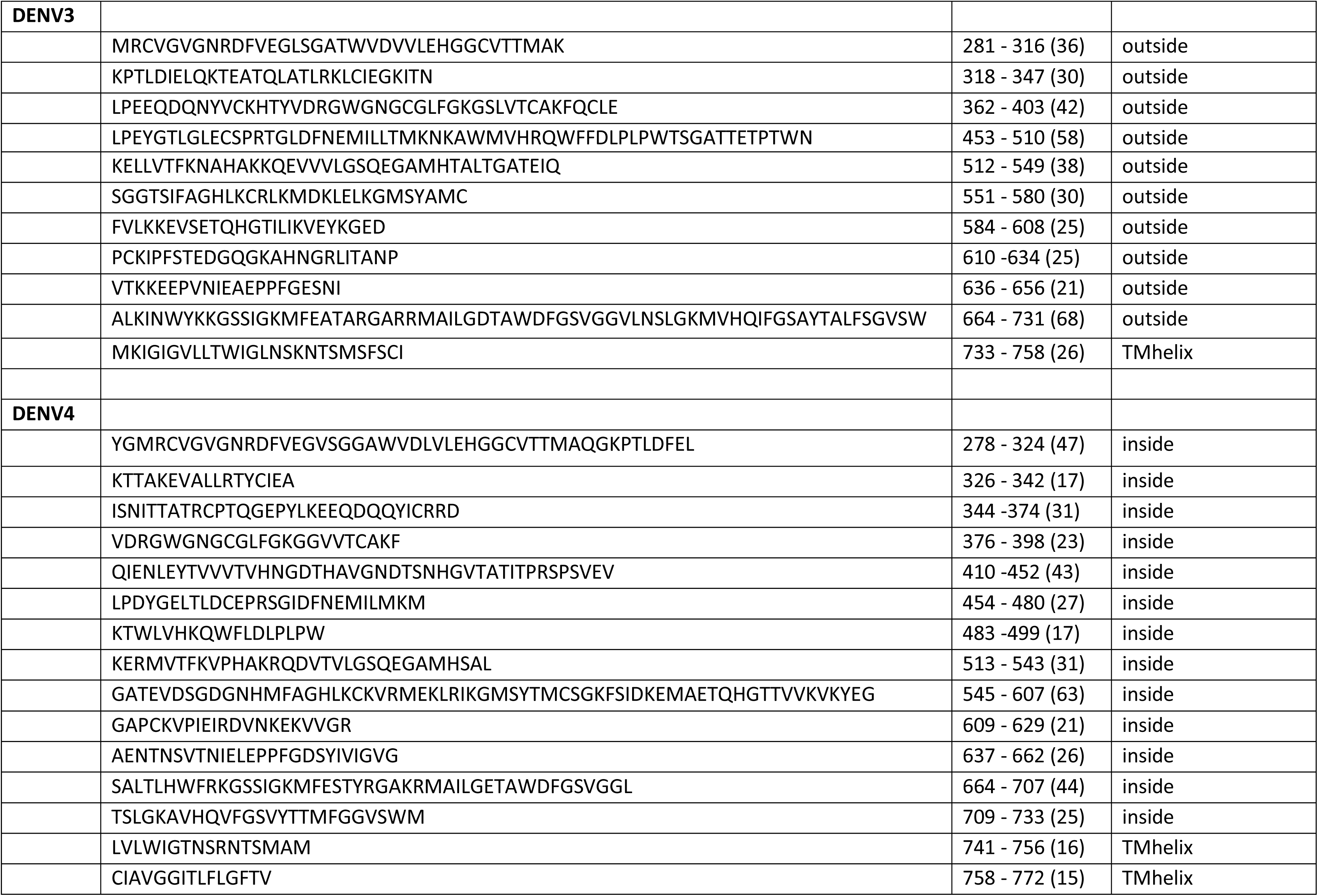
Conserved sequences in the envelop protein of four serotypes of dengue virus.

### Identification of MHC-I Binding T Cells Epitope and Selection from Conserved Domains

The IEDB prediction tools score/ rank different peptides distinctively in different analyses. Such results must be sorted and analysed with an integrated approach to epitope selection. Therefore, we chose a stepwise approach to select epitopes for downstream prediction analyses. Since dengue is most prevalent in humans, only the scores of human MHC-I alleles were used to shortlist the predicted epitopes. However, investigators test the vaccine candidates in preclinical animal models, usually laboratory mice. Thus, the predicted epitopes were also doubly screened for mouse MHC-I alleles in the current study.

In the first step, epitopes with PR<1.0 in H2 MHC-I binding prediction were shortlisted that were also under PR<1.0 in HLA MHC-I binding prediction results (Table 2). The selected epitopes present in the conserved domains of the E protein of the respective serotype were taken for the subsequent analyses. As a result, MHC Class I binding prediction returned 27 predicted epitopes for DENV1 (out of 987 epitopes), 28 epitopes for DENV2 (out of 1180 epitopes), 25 epitopes for DENV3 (out of 964 epitopes) and 41 epitopes for DENV4 (out of 1038 epitopes).

**Table 2:**
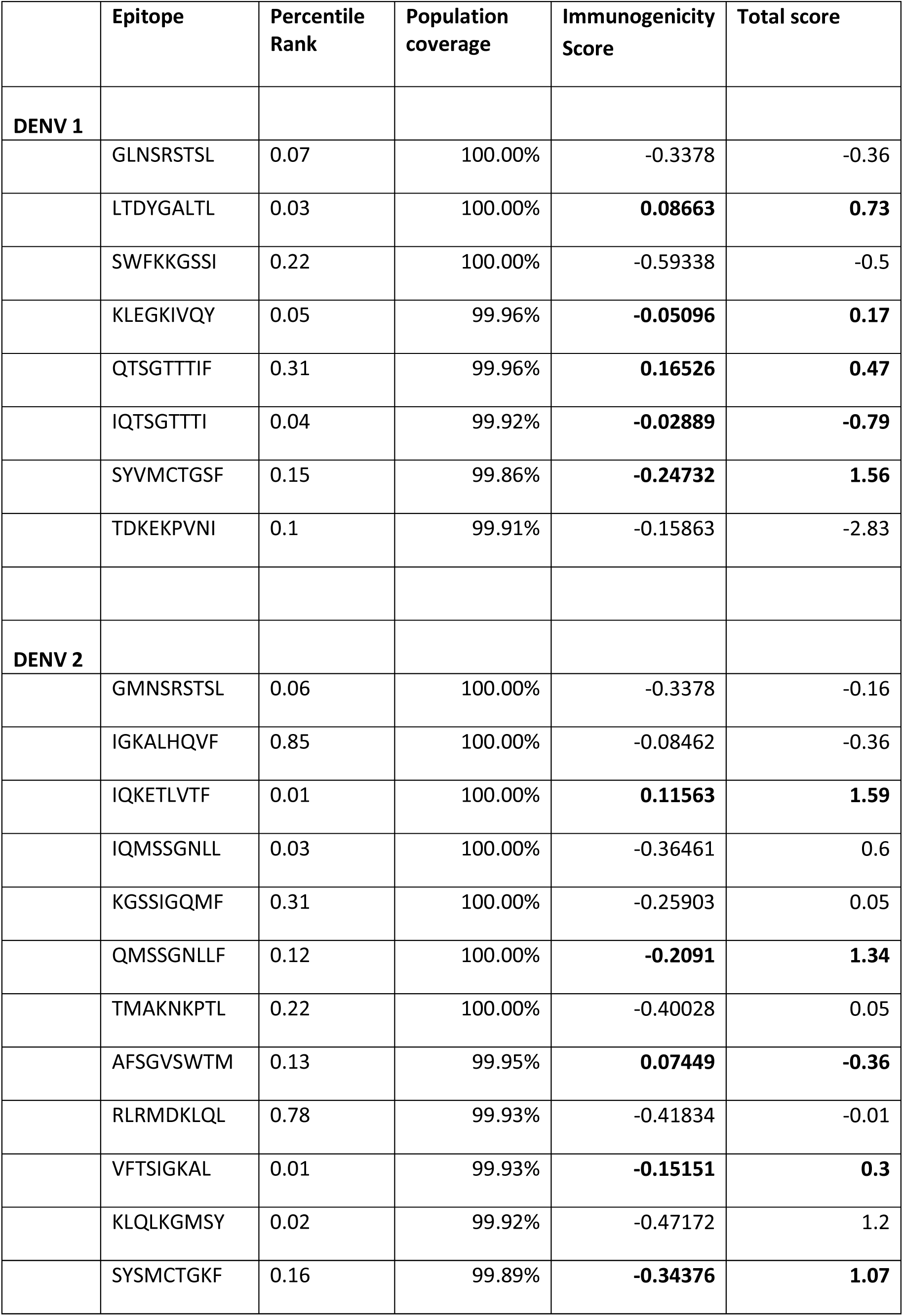
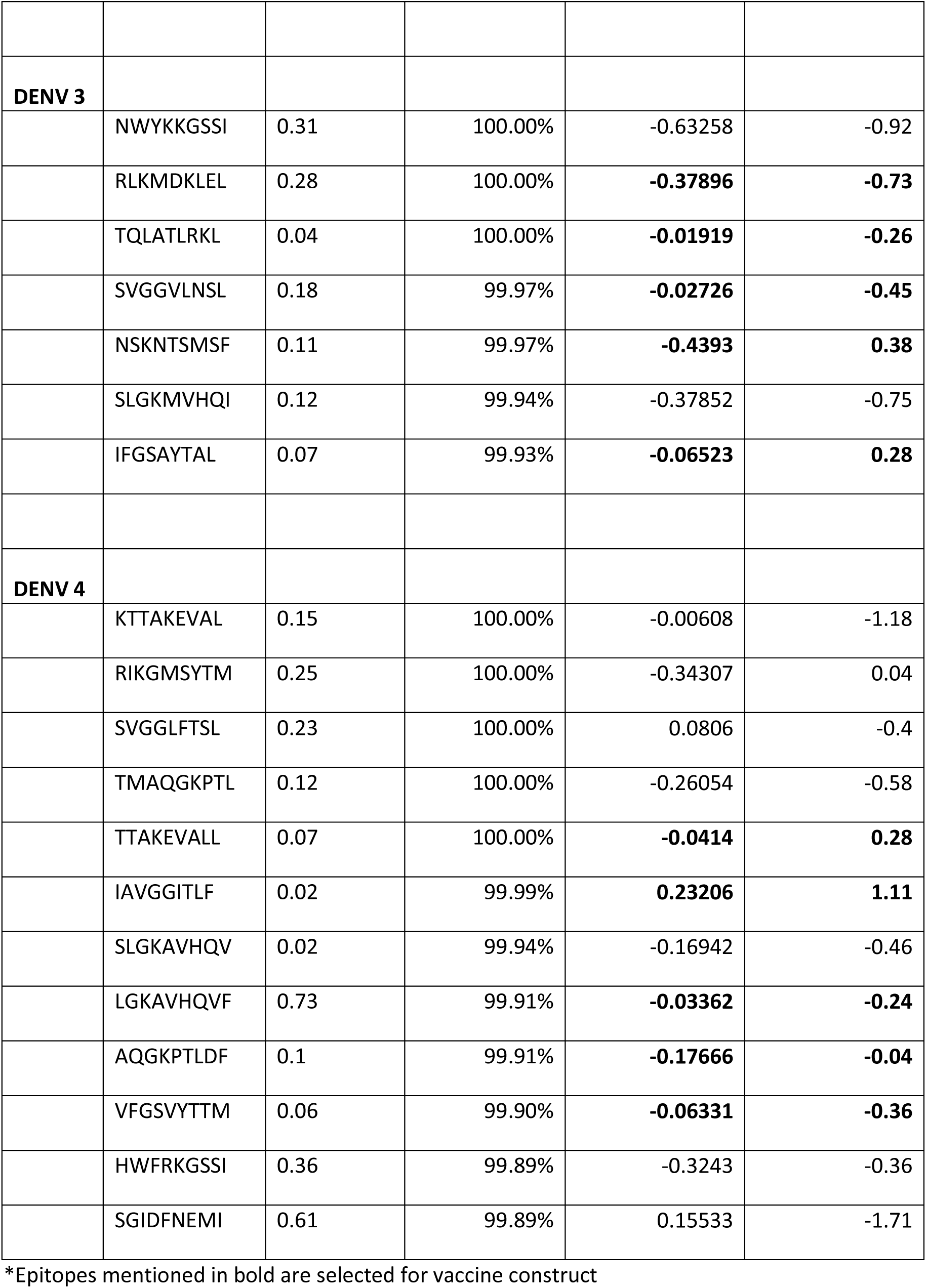
Binding scores, population coverage and immunogenicity score of T cell epitopes selected for each serotype of dengue virus.

Further, a combined prediction was performed for proteasomal processing, TAP transport, and MHC-I binding. The results show the intrinsic potential of the predicted peptide being a T cell epitope.

### MHC Class I Immunogenicity Prediction and Population Coverage

The T cell recognises the pMHC complex if the peptide in the complex is a self- or non-self-antigen. The IEDB MHC Class I immunogenicity prediction tool computes the ability of a peptide as an immunogen. The shortlisted epitopes were screened through this tool and scored, as shown in Table 2. The score value indicates the probability of eliciting an immune response, the higher the score, the greater the probability of immunogenicity. The population coverage was also predicted for the above epitopes. The optimal sets of epitope/HLA combinations were identified using this tool. There were 8 DENV1 epitopes, 12 DENV2 epitopes, 7 DENV3 epitopes and 12 DENV4 epitopes that showed more than 99% global coverage.

### Generation of vaccine construct and its physicochemical characterization

Five most potent MHC-I T cell epitopes from each of the dengue serotypes were joined by AAY linker. The sequence of epitopes and the linker are shown in figure 1. The vaccine construct had a total of 243 amino acids with a total molecular weight of 25.8 kDa and pI of 9.15 which depicts the basic nature of peptide. The construct was predicted to have an average half-life of 4.4 hours in mammalian system and an aliphatic index of 87.08, which indicates its thermostability. The half-life in yeast and bacterial system was predicted to be more than 20 and 10 hours respectively which favours the expression under *in vitro* conditions.

**Figure 1:**
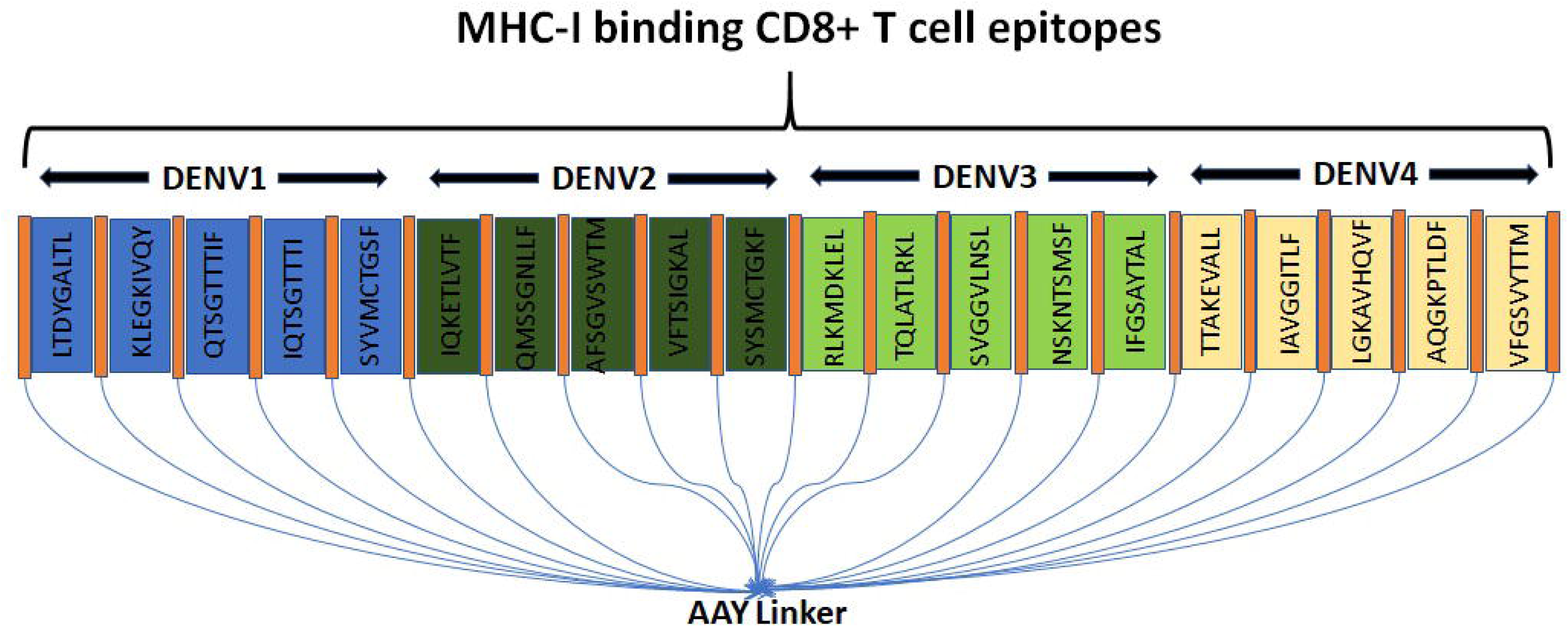
Schematic representation of the chimeric vaccine construct. The predicted CTL epitopes of the four serotypes of dengue virus are linked with AAY linker

### Vaccine construct as a potent antigen and non-allergen

The vaccine construct designed by us was non-allergen as predicted by AllerTOP v2.0. The vaccine construct was also analysed with Vaxijen v2.0 which gave an antigenicity score of 0.4568 at a model threshold of 0.4% showing that the peptide is a probable antigen.

### Structural features of vaccine construct

Analysis of secondary structure of the construct revealed that the peptide had 144 (59.25%), 48 (19.75%) and 51 (20.98%) amino acids in the helix, beta sheets and coil region respectively (Figure 2A). Further the I-TASSER server predicted five models using ten threading templates having the z score between 1.02 to 1.79. We selected the best model having highest C-score of -2.46 for further analysis (Figure 2B). The model was used to generate the Ramachandran plot to assess its accuracy and observed that 93.2 % residues were present in most favoured and additional allowed regions (Figure 2C).

**Figure 2:**
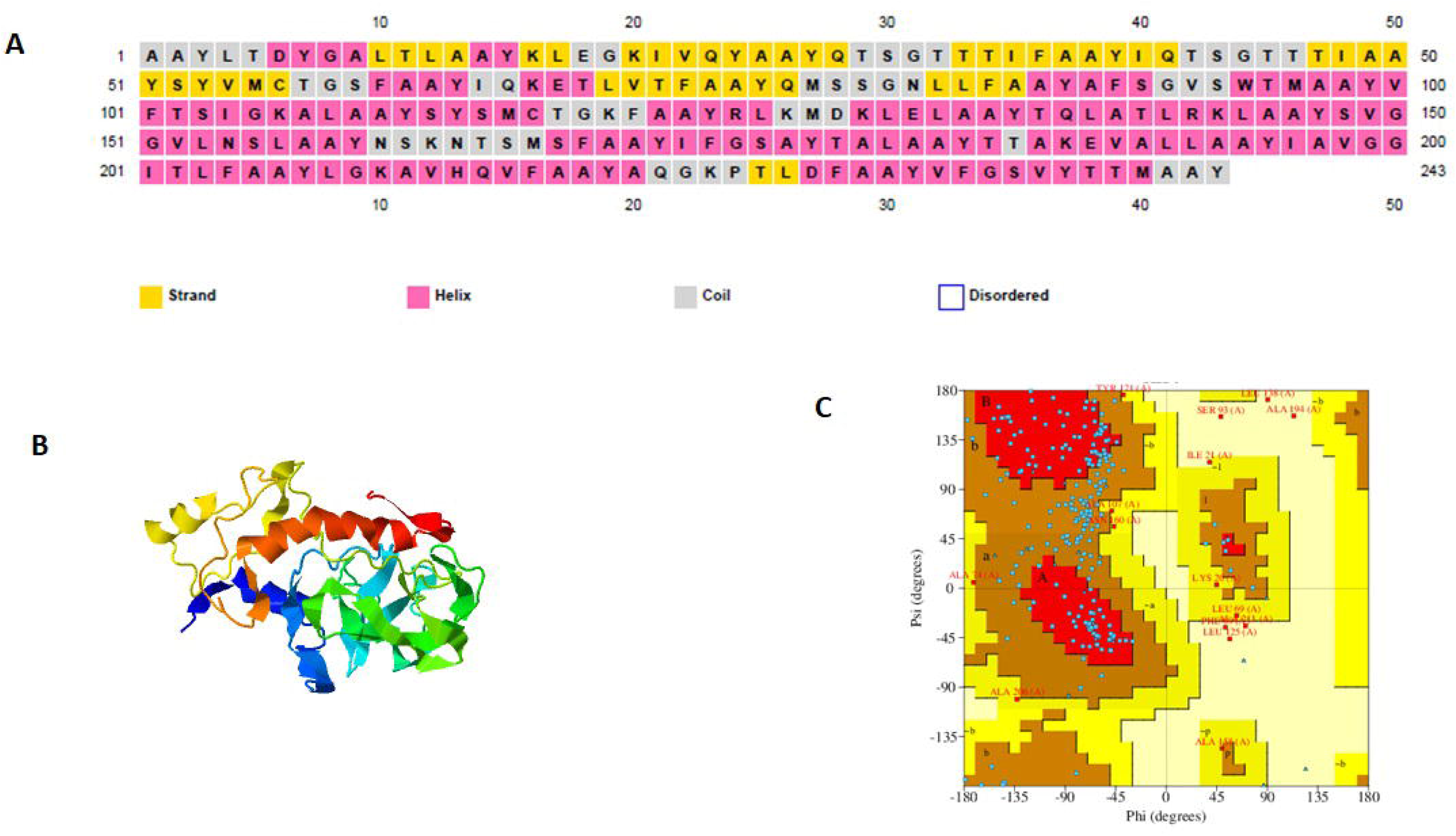
Secondary and tertiary structure of the vaccine construct. (A) Secondary structure of the chimeric construct as predicted by PSIPRED. The structure contains 59.25% helix, 19.75% beta sheet strand and 20.98% coil regions. (B) Best model of tertiary structure of the construct predicted using 1-TASSER. (C) Ramachandran plot analysis for the structure of the construct showing that 93.2% of the residues are present in the most favored and additional allowed regions

### TLR3 interacts with vaccine construct

Docking of TLR3 (PDB ID:2A0Z) with the vaccine construct revealed an effective molecular interaction between them (Figure 3). The LigPlot analysis showed hydrogen bonding and hydrophobic interaction between the TLRs and the vaccine construct.

**Figure 3:**
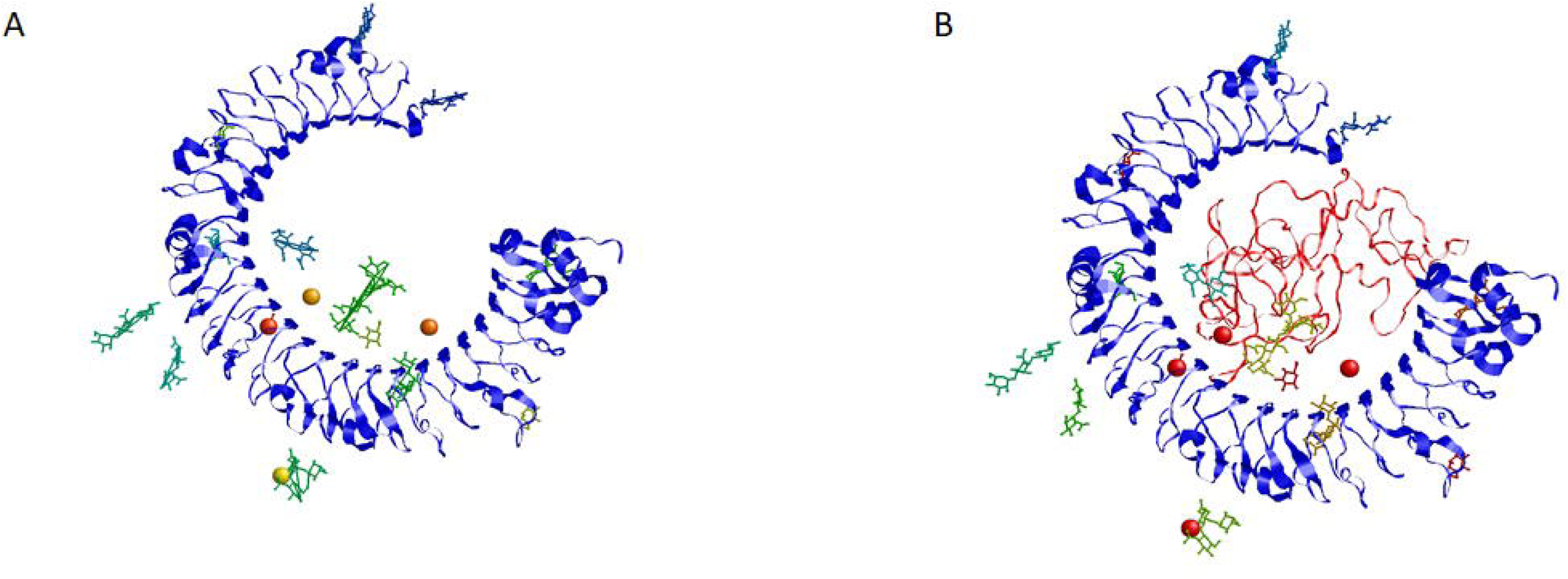
Molecular docking between TLR3 and the vaccine construct using PatchDock online server. A. Structure of TLR3 (PDB ID:2A0Z) B. Docked conformation revealinginteraction between TLR3 and the vaccine construct. Vaccine construct is shown in red.

### In-silico expression analysis of the vaccine construct

Codon optimization was done using Jcat server for maximal expression of the vaccine construct in human cells. The optimized clone encoding the vaccine construct had 729 nucleotides with a GC-content of 66.8% which indicates a high-level expression. The CAI value of >0.95 also indicates the high expression potential of vaccine construct in human cells.

### Computational immune simulation reveals effective cytotoxic T cells and Th1 cytokine response

C-ImmSim online server was used to predict the immune response towards the vaccine construct when three injections were given at four weeks interval. Immune simulation revealed that the cytotoxic T cells are activated exponentially within 5 days of injection which persists up to more than 300 days. The Th1 cytokine response was also predicted with IFNγ being the most prevalent cytokine followed by Interleukin2 (IL2) after the third immunization (Figure 4).

**Figure 4:**
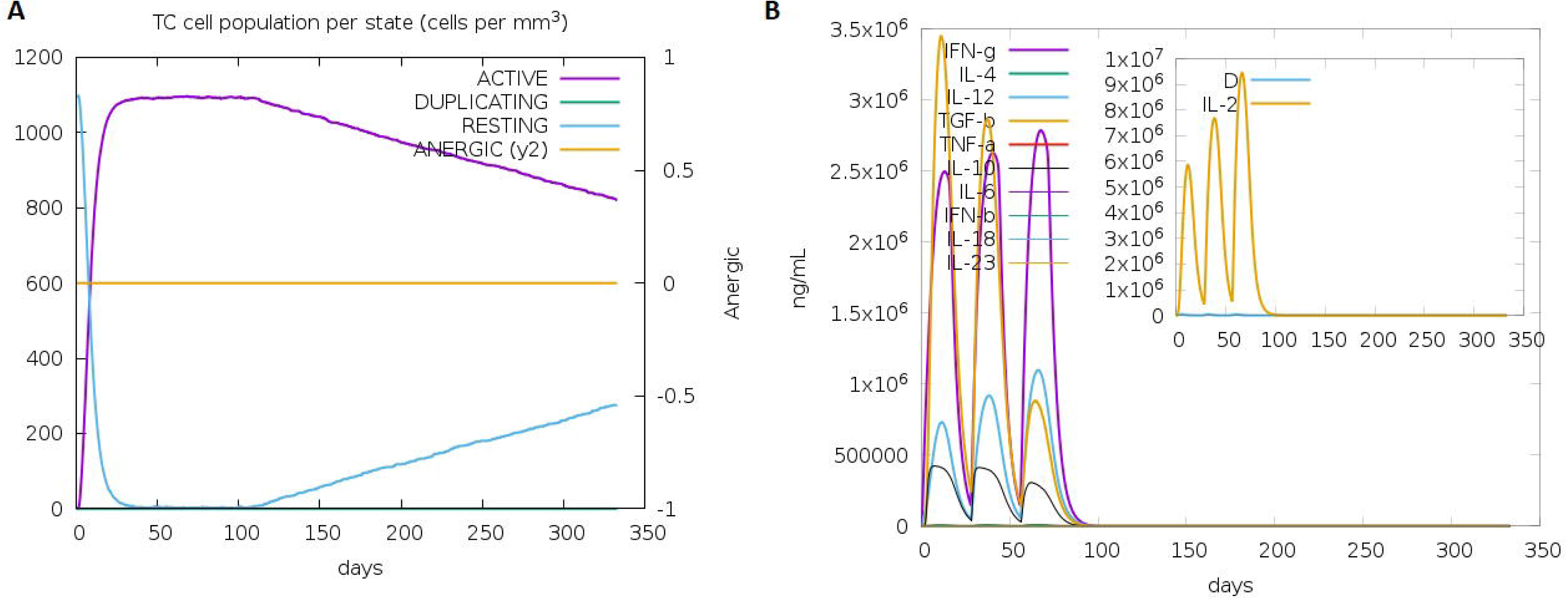
Simulation of immune response usingC-lmmSim online server after three injections of the vaccine construct given at four weeks interval. (A) Evolution of Cytotoxic T cell population. (B) Cytokine and interleukin production and persistence after each injection.

## Discussion

The modern world has seen the emergence of viral infections which have adventitiously appeared in regions other than the tropics and subtropics. Many countries like the Americas, Africa, the Middle East, Asia, and the Pacific Islands have experienced varying magnitudes of dengue outbreaks (CDC, 2023). While most dengue cases are asymptomatic, they can feed mosquitoes for DENV transmissions to high-risk human communities of infants, young adults, immune-compromised or older people (Duong et al., 2015). Besides this, severe complications of dengue cases devoid of prompt diagnosis and medical care may alarmingly exceed the fatality rates above 20% (Hentzy et al., 2013). Although there are a few dengue vaccines in the pipeline, only one tetravalent live attenuated vaccine (Dengvaxia; Sanofi Pasteur) is approved by the regulatory authorities in nearly 20 countries and has been shown efficacious only in seropositive individuals (Amorim et al., 2022). Therefore, safer and more efficient vaccine strategies may fill the vacuum in integrated dengue prevention and control measures.

The advent of machine learning platforms has resolved the laborious tasks of construction and assessment of protein-based vaccine designs. In this study, we have attempted to design a chimeric protein by linking the epitopic peptides, which could be examined for their efficacy in eliciting a cell-mediated immune response. It is well established that the cytotoxic CD8^+^ T lymphocytes (CTL) restrict dengue viremia or at least limit disease severity by recognising and killing infected cells (Tian et al., 2019). Training our immune system using dengue-specific MHC Class I binding T cell epitopes may build strong CTL immunity against the four DENV serotypes (Testa et al., 2012). The E protein of DENV showcases a vital role in the virus attachment and later in the assembly of viral particles and the formation of mature virions (Perera et al., 2008). CTL responses are predicted to be more inclined towards proteins which have high abundance in native infection and are evolutionarily conserved (Bilderbeek et al., 2022; Khan et al., 2008). The highly conserved domains of the E protein of DENV were identified for each serotype in this study and assessed for their immunogenicity. Immunogenicity is not only dependent on the T cell epitope but also on the frequency of the class I HLA allele in the population (Weiskopf et al., 2013). Hence, population coverage prediction of the compiled epitopes was also assessed, which covered more than 99% percent of the population for each serotype. However, as given in Table 2, the individual epitopes estimated a varying coverage. Therefore, only the epitopes showing more than 50 percent population coverage and relatively higher immunogenicity scores were selected for docking analysis.

Our vaccine construct has twenty T cell epitopes and is composed of 243 amino acids. The epitopes were linked to each other with AAY linker. The AAY linker is known to be the cleavage site for proteasomes in mammalian system which helps in effective separation of epitopes within the cells without any junctional immunogenicity (Bhatnagar et al., 2020). As described previously by Yang et al., the linker is expected to enhance the immunogenicity of our chimeric peptides (Yang et al., 2015). In addition to be immunogenic, a vaccine candidate should also be a good antigen and should not be a potential allergen. Therefore, we determined the antigenic and allergenic characteristic of the vaccine construct using AllerTOP v2.0 and VaxiGen v2.0 tool whereby we determined that the construct is an antigen but not an allergen. Structure of a vaccine candidate determines its stability under physiological conditions and its recognition by the immune components of recipient body. Our prediction reveals that 59.25 % of the chimeric peptide is in the helix region, 19.75% is present as beta sheet and 20.98% is present in the coil region. The tertiary structure determined by I-TASSER shows that 93.2% of the peptide is in the most favoured region or in the additional allowed regions which signifies that the structural quality of the chimeric peptide is acceptable as a vaccine candidate. We also determined whether the peptide can be recognized by TLRs, and our docking analysis revealed an effective binding affinity between them.

Further, the immune simulation with C-ImmSim server showed the significant activation of CTLs within 5 days of injection which persists for up to 300 days along with Th1 cytokine response.

In conclusion, the conserved epitope selection for each DENV serotype in this paper is demonstrated for the first time. Although the immunoinformatics approaches used in the current study can provide highly accurate predictions. However, these epitopes should be assessed *in vitro* and *in vivo* for credibility as therapeutic peptide candidates. Individual peptides of each DENV serotype should be tested in CD8 T cell expansion assays and analysed for the peptide-specific interferon-gamma response in the effector cells. The vaccine construct further needs to be tested in experimental animal models to check its efficacy against all four serotypes of dengue virus.

## Acknowledgement

This work was supported by Indian Council of Medical Research (ICMR), New Delhi vide letter No. VIR/Fellowship/35/2019-ECD-I

## Author Contribution

N.M.: study design, execution, acquisition of data, analysis and interpretation of results, and manuscript preparation; P.K.: conception, supervision, review and editing original draft, analysis and interpretation of results, project administration; M.K.: supervision, project administration, formal analysis.

